# Sensorimotor adaptation reveals systematic biases in 3D perception

**DOI:** 10.1101/2025.01.22.634281

**Authors:** Chaeeun Lim, Dhanraj Vishwanath, Fulvio Domini

## Abstract

The existence of biases in visual perception and their impact on visually guided actions has long been a fundamental yet unresolved question. Evidence revealing perceptual or visuomotor biases has typically been disregarded because such biases in spatial judgments can often be attributed to experimental measurement confounds. To resolve this controversy, we leveraged the visuomotor system’s adaptation mechanism — triggered only by a discrepancy between visual estimates and sensory feedback — to directly indicate whether systematic errors in perceptual and visuomotor spatial judgments exist. To resolve this controversy, we leveraged the adaptive mechanisms of the visuomotor system to directly reveal whether systematic biases or errors in perceptual and visuomotor spatial judgments exist. In a within-subject study (N=24), participants grasped a virtual 3D object with varying numbers of depth cues (single vs. multiple) while receiving haptic feedback. The resulting visuomotor adaptations and aftereffects demonstrated that the planned grip size, determined by the visually perceived depth of the object, was consistently overestimated. This overestimation intensified when multiple cues were present, despite no actual change in physical depth. These findings conclusively confirm the presence of inherent biases in visual estimates for both perception and action, and highlight the potential use of visuomotor adaptation as a novel tool for understanding perceptual biases.

## Introduction

A fundamental question in sensory perception research concerns the nature of biases in the perception of spatial properties (e.g., direction, distance or depth) and how these biases impact visually guided actions. Previous studies have shown systematic biases in depth estimation for both perception^1–6^ and action^7–9^. One common finding is that the estimated depth of an object is not constant across distances. Specifically, object depth appears greater than that specified by disparity information within reaching distances, but is underestimated at far distances^10–16^. Figure 1A illustrates a disparity only stimulus of the sort in which such overestimation bias at near distances is observed. Another spatial bias repeatedly observed in the literature is that depth estimation varies depending on the number of depth cues involved: with multiple depth cues, objects appear deeper than when only one cue is available, despite no difference in rendered depth. This has been observed in both perceptual^17,18^ and visuomotor tasks^19^. Figure 1B illustrates an object demonstrating such overestimation.

**Figure 1.**
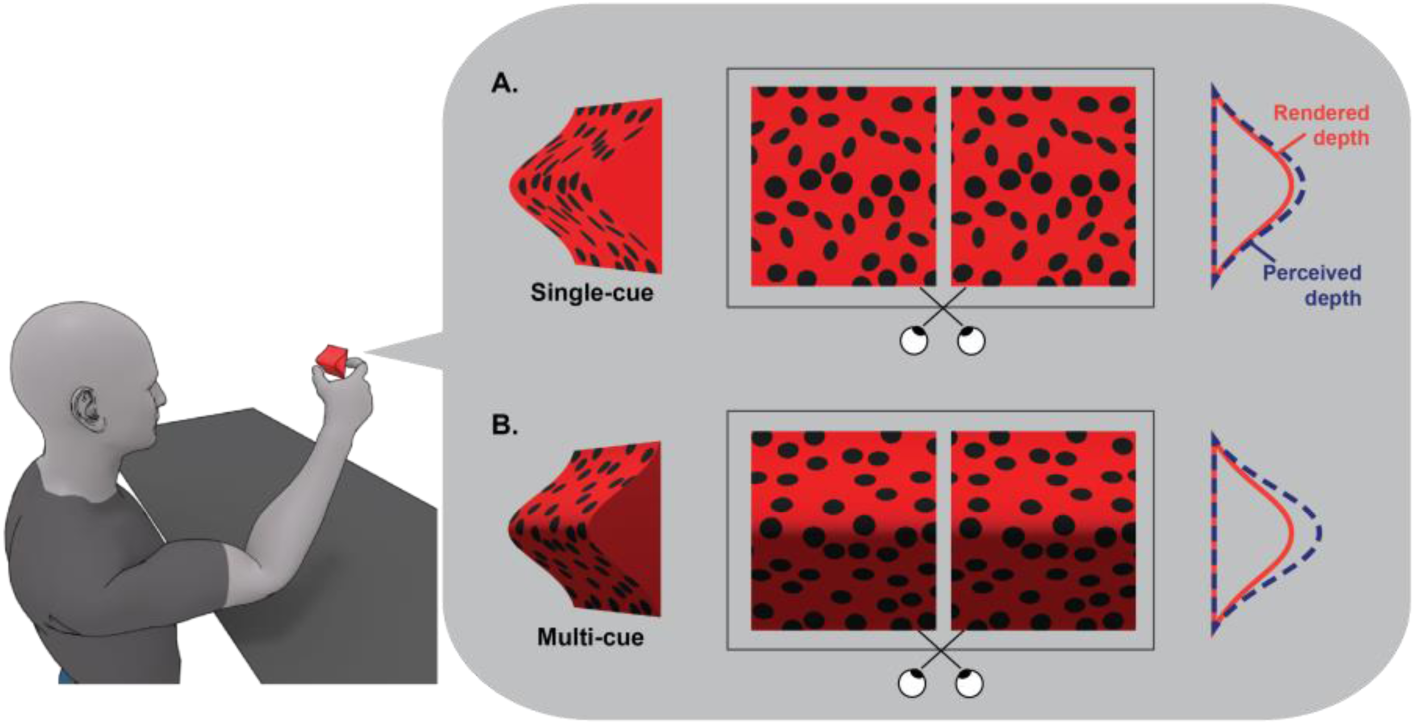
An example of a task where the observer perceives and grasps a virtual object specified by different cues. The left-most images depict 3D stimuli at a 45° orientation for illustration purposes. The middle images are the stereoscopic version, observable through cross-fusing. The right-most diagrams show the true rendered depth profile (red) and the pattern of perceived depth profiles for such stimuli (dark blue). A. An example of a disparity only (single cue) stimulus where an overestimation of depth (dark blue) can be observed in comparison to the rendered ground truth (red). B. A multi-cue stimulus (disparity, texture gradient and shading) which has been shown to generate even greater overestimation of depth (dark blue) in comparison to the disparity only stimulus, despite their same rendered depth.

Despite extensive evidence of bias in depth estimation, it remains unresolved whether our depth perception is indeed biased and leads to errors in subsequent actions. Some propose that the human visual system can accurately perceive 3D structures under natural viewing conditions^20–30^, where bias found in previous studies is typically attributed to artifacts caused by virtual environments^16,31^. For instance, this perspective explains that the multi-cue stimulus may appear deeper than the single-cue stimulus (Figure 1) because the latter is underestimated. In virtual environments, depth cues such as accommodation, defocus blur, and disparity typically indicate the zero depth of the flat screen, often called ‘flatness cues’^16,31,32^. Integrating these cues with other cues specifying non-zero depth leads to an inaccurate underestimation of object depth. However, as more depth cues are added, the influence of flatness cues diminishes, making the multi-cue stimulus depth appear greater due to less underestimation.

Another perspective argues that although perception can be biased, this bias does not impact subsequent actions^33–35^. According to this perspective, in the example shown in Figure 1, even if the multi-cue stimulus perceptually appears deeper than the single-cue stimulus, it would not lead to a similarly overestimated grip during a grasping action. However, whether the sensory bias consistently affects both perception and action remains unclear, as previous studies suggest mixed results. Some findings showed that visual illusions distort perceptual judgments but not actions like grasping^36–38^, while other studies observed comparable biases in both perception and action^39–41^.

The challenge in elucidating the existence of sensory bias lies in the difficulty of measuring it in absolute terms. Bias is conventionally defined as the systematic discrepancy between physical measurements (i.e., ground truth) and the visual estimates from the observer’s perceptual judgments or visuomotor estimates (e.g., grip size). However, visual estimates are typically inferred from a participant’s performance on tasks like stimulus comparison, probe matching, or manual size estimation, which themselves can introduce task-related biases. This confounds the true measurement of the sensory estimate. For example, the most common method, the two-interval forced-choice (2-IFC) task, measures the point of subjective equality (PSE) by comparing a stimulus depth to a reference^42–44^. The PSE reflects the point where the stimulus is perceived as equivalent to the reference, but this measure is relative and thus susceptible to biases in the perceived depth of the reference. Similarly, in a probe adjustment task, participants match a 2D probe to the estimated profile of a 3D target stimulus^45–48^. However, this requires mental rotation, potentially introducing bias and individual differences irrelevant to depth estimation itself. More direct measures like a manual estimation task ask participants to adjust their thumb-to-index finger distance to indicate perceived depth or size^17,49^. However, reliance on proprioception also introduces potential biases due to individual differences in proprioceptive sensitivity to the distance between digits, especially in the absence of visual feedback of the hand^50^.

Our aim was to conclusively determine if bias in visual depth estimation exists and is significant. To circumvent the challenges of measuring the sensory estimate with perceptual judgment tasks, we propose a novel approach that capitalizes on the visuomotor system’s existing ability to detect perceptual errors. Instead of trying to directly measure the estimate, we measure visuomotor adaptation, a corrective mechanism used by the visuomotor system to adjust for discrepancies between planned actions and desired outcomes. Consider the previously mentioned bias where stimuli specified by multiple depth cues often appear deeper than their actual depth within reach space (Figure 2). If such a bias exists, when grasping such an object, the visuomotor system should initially plan a larger grip due to the overestimated depth. Upon receiving haptic feedback indicating the physical depth, it should rapidly adjust the mappings between visual estimates and planned grip to accurately match the physical depth in subsequent trials. Thus, given the same visual information, the visuomotor system now plans for a smaller grip.

**Figure 2.**
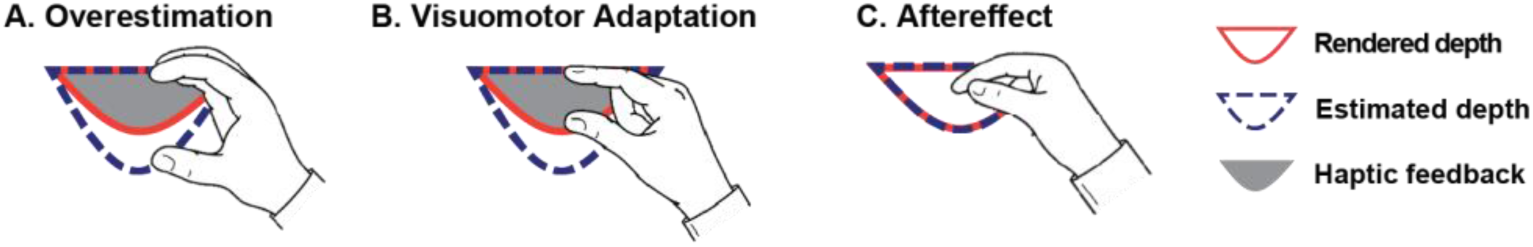
An example of the visuomotor adaptation process in a grasping task. The red circular segment represents the true rendered depth, the dark blue dashed segment represents estimated depth, and the grey segment represents haptic feedback. (A) If the stimulus appears deeper than its rendered depth, planned grip will be overestimated. (B) However, haptic feedback signaling the true rendered depth triggers the visuomotor system to adapt to make a smaller grip relative to the estimated depth. (C) If the stimulus is switched to one having the same rendered and estimated depths, the planned grip would be smaller than before as an aftereffect.

Note that the visuomotor system initiates this adaptation mechanism only when it recognizes a discrepancy between visual estimates and haptic feedback, indicating the presence of sensory bias. Thus, the occurrence of visuomotor adaptation implies that bias is present and causes significant error in visuomotor estimates. Furthermore, the direction of the bias indicates the sign of the error in visual estimates. Leveraging these characteristics of visuomotor adaptation, the present study aims to incontrovertibly establish whether biases in perceptual depth estimation exist even when signals unambiguously specify the true depth.

We designed a grasping task where participants repeatedly grasped 3D objects to specifically test the effect of depth overestimation in multi-cue stimuli compared to single-cue stimuli. While the stimulus was presented virtually, participants grasped a real physical object at the end of each trial that was aligned with the stimulus, providing haptic feedback about the stimulus depth (Figure 3). Maximum Grip Aperture (MGA) was used to infer the planned grip size^51–53^. The task consisted of four stages based on stimulus type (Figure 4). In Stage 1, we presented a single-cue stimulus defined solely by binocular disparity information. We were agnostic about whether disparity alone leads to overestimation, underestimation, or veridical perception of depth. For simplicity, we predicted that the planned grip (yellow) would initially be around the rendered depth and that specified by haptic feedback (grey). Regardless of initial overestimation or underestimation, experiencing haptic feedback would eventually bring the planned grip to converge to the haptically specified depth through visuomotor adaptation, which served as a baseline for the following Stages.

**Figure 3.**
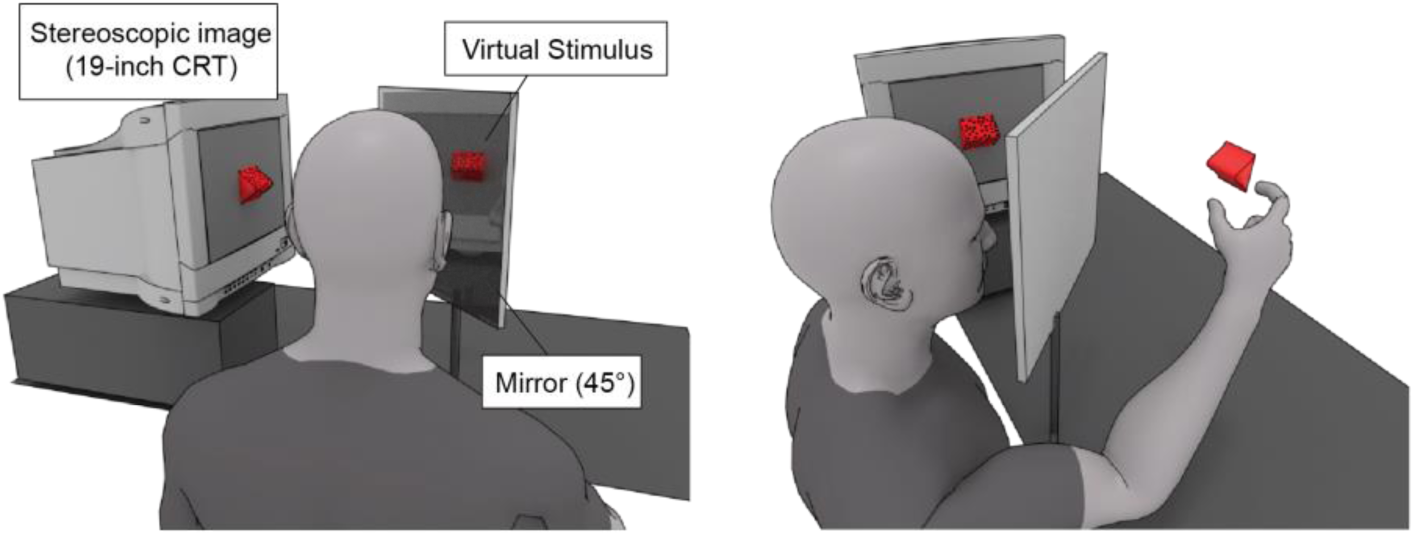
The illustration of the experimental setup. (left) The stereoscopic images were reflected in a mirror. When viewed through shutter glasses, these images appeared as a 3-dimensional object floating in front of the participants. (right) During grasping, participants reached toward the virtual stimulus to grasp it, but they actually grasped the physical object aligned with the virtual stimulus.

**Figure 4.**
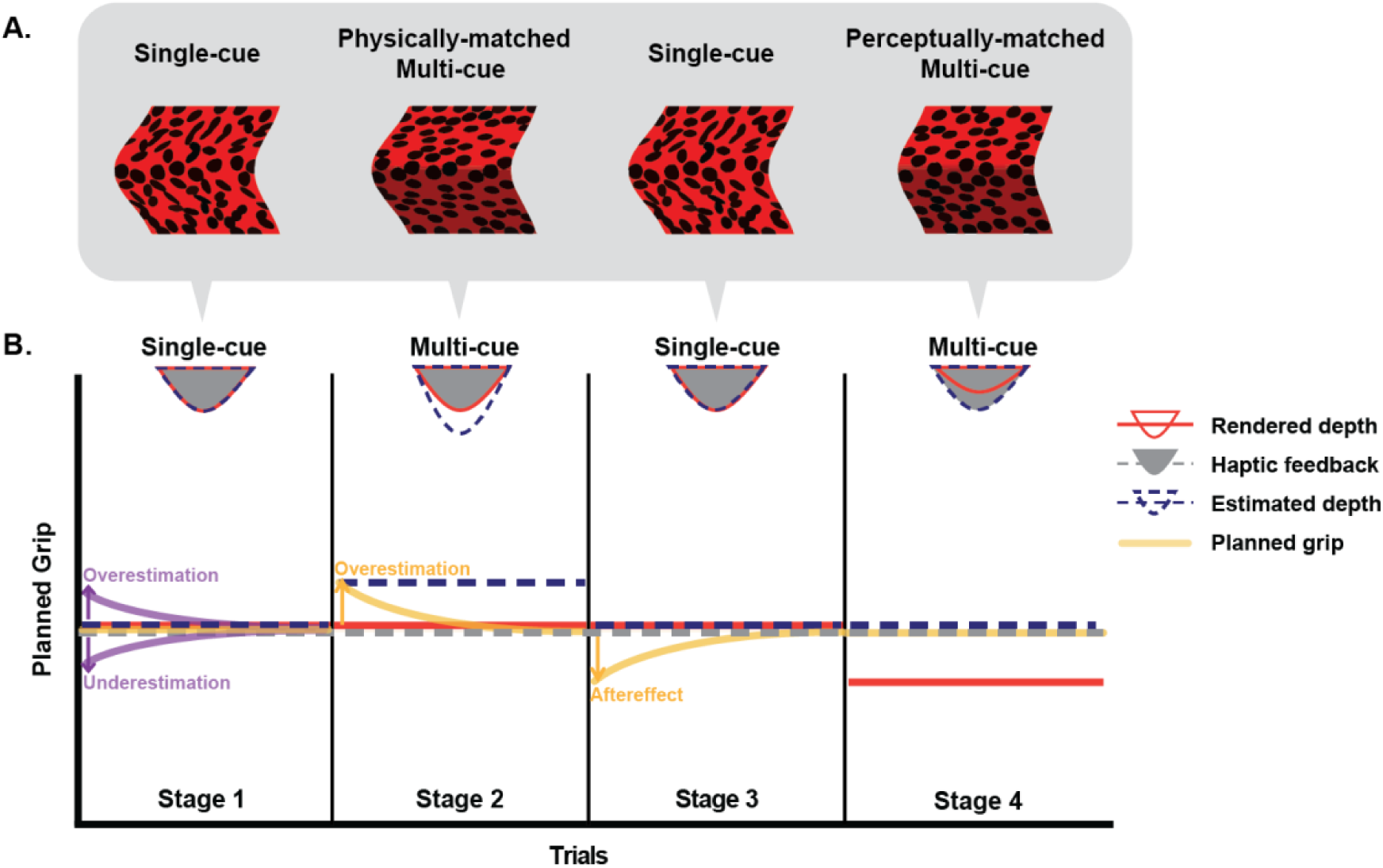
A. The stimuli used in different stages of grasping tasks. B. Each stimulus’ rendered depth (red) and estimated depth (dark blue) are illustrated with semi-circular diagram on top of the schematic plot showing four stages of the grasping task. The haptic feedback (grey) specified the rendered depth of the stimuli in Stages 1, 2, and 3, but it was larger than the rendered depth of the perceptually-matched multi-cue stimulus in Stage 4. In Stage 1, depth from binocular disparity alone could yield a veridical estimate (aligning with the rendered depth and haptically-specified depth) or it could be either overestimated (due to inherent bias in depth perception) or underestimated (due to flatness cues). The over- and underestimation are shown in purple. Irrespective of initial over- or underestimation, the planned grip apertures will eventually converge to haptically specified depth at the end of Stage 1 through visuomotor adaptation. In Stage 2, additional depth cues are added to the stimulus while keeping the rendered depth constant (i.e., the multi-cue stimulus depth is physically matched to that of the single-cue stimulus). If previously reported depth overestimation for multi-cue compared to single-cue stimuli exists, it should cause a sudden increase in planned grip at the beginning of Stage 2. The planned grip should decrease and converge with the haptically-specified depth (visuomotor adaptation). If multi-cue stimulus overestimation is observed in Stage 2, then at the beginning of Stage 3, where the single cue stimulus is reintroduced, there should be a sudden decrease in planned grip (aftereffect). In Stage 4, where a multi-cue stimulus matched in perceived depth to the single cue stimulus (i.e., perceptually matched), no change in planned grip is expected because the perceptually estimated depth in Stage 3 and 4 should be the same.

There are two possible scenarios in the initial phase of Stage 1. If the depth from binocular disparity is overestimated within reaching distance, as observed in some literature^11,14,16^, and this bias affects motor planning, initial grips should reflect overestimation (purple, above). Conversely, if the bias in depth perception is due to artifacts from virtual environments, such as flatness cues specifying the monitor screen, the initial planned grip should be underestimated (purple, below). The latter bias can be explained on the basis that the non-zero depth from binocular disparity would be combined with conflicting flatness cues specifying the zero depth of the screen, resulting in a final depth estimate smaller than the depth originally rendered with the disparity^16,31^. In either case, haptic feedback indicating the discrepancy between the planned grip and the rendered depth of the virtual stimulus should prompt visuomotor adaptation. However, the direction of the adaptation will distinguish whether the initial planned grip is overestimated or underestimated.

In Stage 2, the same stimulus depth as in the previous stage (red) was specified by multiple cues including disparity, texture gradient, and shading. Thus, the multi-cue stimulus depth was physically matched to the single-cue stimulus depth (‘physicallymatched multi-cue’). However, prior works have demonstrated that the depth of the multi-cue stimulus is overestimated compared to the single-cue stimulus^18,19^. If this finding is valid, the multi-cue stimulus in Stage 2 should be perceived as deeper (dark blue) than the disparity-only stimulus previously grasped in Stage 1, resulting in an immediate increase in the planned grip (yellow). Subsequently, the discrepancy between haptic feedback (grey) and planned grip should again induce adaptation and an accompanying reduction in planned grip over subsequent trials.

In Stage 3, reintroducing the single-cue stimulus allowed us to assess any residual adaptation effects from Stage 2. If the single-cue stimulus indeed appears shallower than the multi-cue stimulus, the planned grip (yellow) should initially be underestimated due to the decrease in estimated depth (dark blue) and the negative aftereffect from the previous phase, followed by an increase due to visuomotor adaptation from haptic feedback (grey).

Finally, in Stage 4, a multi-cue stimulus was reintroduced, but with its rendered depth (red) adjusted to match the perceived depth (dark blue) of the single-cue stimulus in the previous stage. To personalize this adjustment for each participant, participants first completed a perceptual matching task before the main grasping task. In this adjustment task, they adjusted the rendered depth of the multi-cue stimulus until it appeared as deep as the single-cue stimulus, creating perceptually matched pairs of single-cue and multi-cue stimuli. The haptic feedback (grey) matched the rendered depth of the single-cue stimulus (i.e., smaller than the estimated depth of the multi-cue stimulus). Since there is no change in either the estimated depth or the haptic feedback from Stage 3, we expect no changes in planned grip (yellow).

The planned grip expectations outlined in Figure 4 are based on the assumption that actions are planned using the perceptual estimate of depth, which may be biased by different factors. Alternatively, if the visual system can accurately derive estimates for guiding action^33–35^, the estimated depth (dark blue) should always align with the rendered depth (red, Figure 5). Thus, the planned grip (yellow) would be constant throughout Stages 1, 2, and 3, as there is no change in both rendered depth and haptic feedback (grey). It would abruptly decrease only in Stage 4 when the rendered and thus estimated depths are smaller than those of the stimuli presented in the preceding Stage 3. Then, the planned grip would converge back to the haptically-specified depth through visuomotor adaptation.

**Figure 5.**
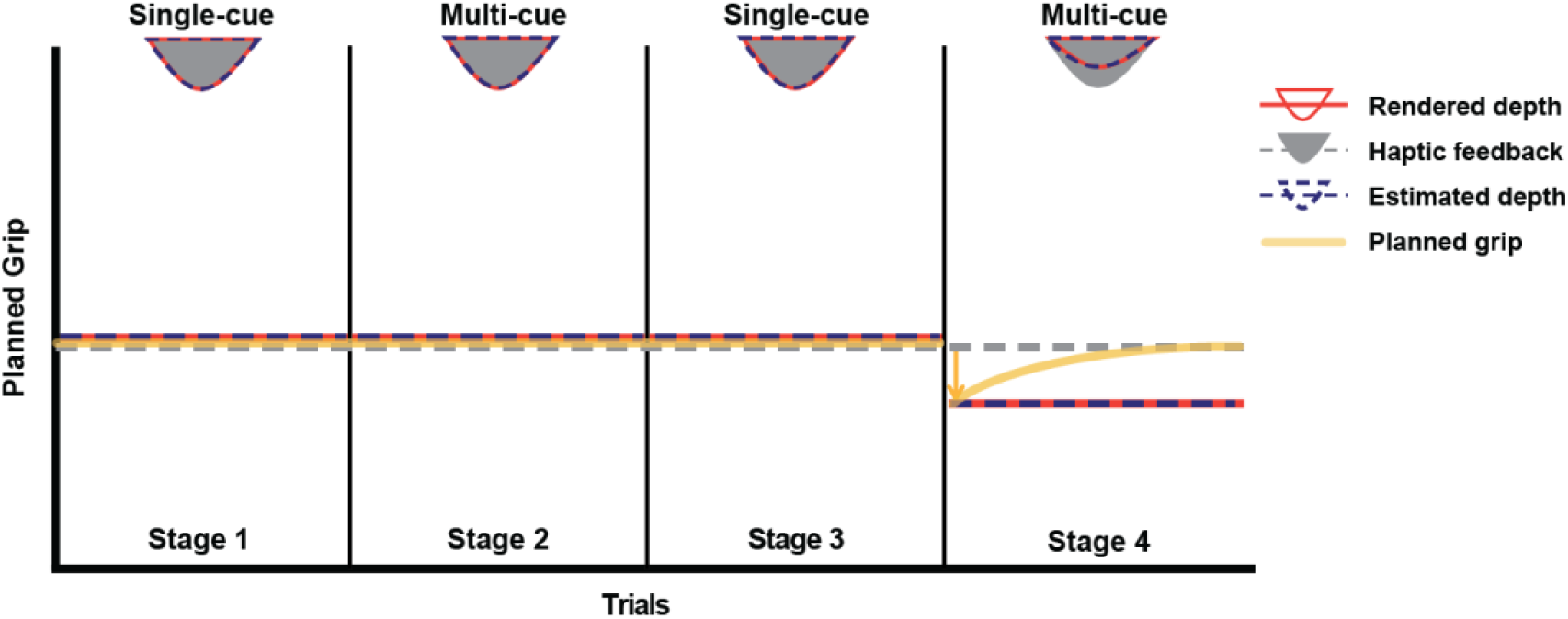
The estimated depth and the planned grip predicted in the case where the visual system can accurately estimate rendered depth for guiding action, regardless of how many depth cues are involved. During the initial three stages, the true rendered depth (red), estimated depth (dark blue), and haptic feedback (grey) are expected to align with each other, resulting in a constant planned grip (yellow) throughout. However, in Stage 4, a decrease in rendered depth, and consequently, in estimated depth, would result in an abrupt decline in the planned grip. Subsequently, the grip is expected to converge back to baseline levels after receiving haptic feedback (visuomotor adaptation).

## Results

### Perceptual Matching Task

Participants first completed a perceptual matching task to create a personalized set of multi-cue stimuli that matched the depth of the single-cue stimuli. They adjusted the depth of a multi-cue stimulus to match a single-cue stimulus with a fixed depth of 27 mm, 30 mm, or 33 mm. Participants adjusted the depth of the multi-cue stimulus (disparity + texture + shading) to be smaller (approximately 3.4 mm on average) than that of the single-cue stimulus (disparity only) to obtain a perceptual match, *t*(23)= 8.13*, p < .*001, (Figure 6). That is, the multi-cue stimuli were perceived deeper than the single-cue stimuli. This increase in perceived depth was significant for each tested depth magnitude of the multi-cue stimulus: the adjusted depth was smaller than the single-cue stimulus depth by approximately 2.7 mm for the small, 3.4 mm for the medium, 4.0 mm for the large depth magnitudes (*t*(23)= 5.94*, p < .*001*,t*(23)= 7.56*, p < .*001, and *t*(23)= 9.33*, p < .*001, respectively). Consistent with previous findings, this result corroborates evidence that providing more depth cues increases the perceived depth.

**Figure 6.**
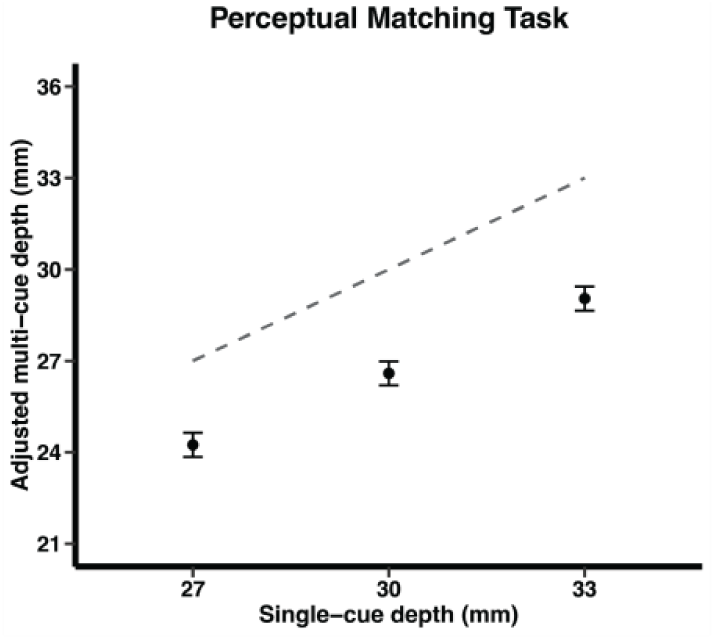
Perceptual matching task result. The depths of the single-cue stimuli (standard stimuli) are plotted on the x-axis, and the average depths of the multi-cue stimuli (probe stimuli) that participants adjusted are plotted on the y-axis. The dashed line denotes an identity line. The depths of multi-cue stimuli were adjusted to be smaller than the single-cue stimuli depths, which indicates that they appeared deeper than the single-cue stimuli. The error bar denotes 95% within-subject confidence intervals^54^.

### Grasping Task

We used MGA to infer the planned grip size^51–53^. Three different depth magnitudes were presented in a pseudo-random order, ensuring no repetition until all three depths were shown. We defined a set of three trials with different depths as a bin and averaged the MGA for each bin. We examined how MGAs vary with the presentation of different stimulus types and the experience of haptic feedback (Figure 7). To assess visuomotor adaptation, we applied either an exponential or linear model^55,56^—whichever provided the better fit—and compared it to a constant model indicating no adaptation. In addition, we conducted correlational analyses to test whether the fitted model could predict the observed MGAs.

**Figure 7.**
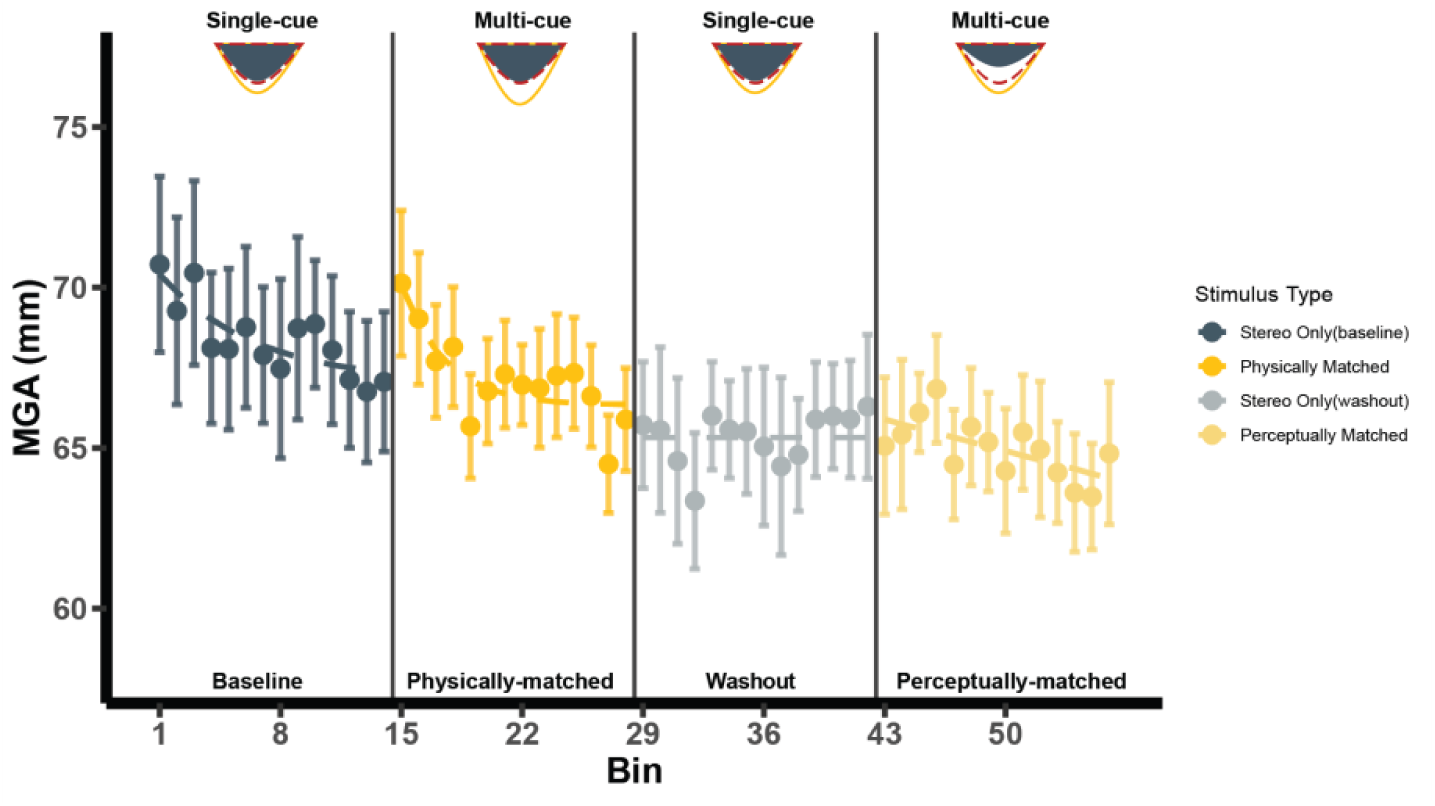
Grasping task results. The bin consists of 3 trials of varying depth magnitudes plotted on the x-axis, while the average MGA of each bin is plotted on the y-axis. The dashed line is the best-fitting model: exponential models for Stages 1 and 2, the constant model for Stage 3, and the linear model for Stage 4. The error bar denotes 95% within-subject confidence intervals^54^.

In Stage 1, the data reveals a trial-to-trial reduction in MGA consistent with visuomotor adaptation that compensates for a perceptual overestimation. The exponential adaptation model yielded a better fit to the data (*AIC* = 36.39) compared to the constant (no change) model (*AIC* = 47.53) and the MGAs predicted by exponential models were significantly correlated with the observed MGAs, *r*(12)= .81*, p < .*001 (Figure 8). This suggests that the visually perceived depth from the disparity cue alone was overestimated, and the visuomotor system detected and corrected its discrepancy from the true depth by using haptic feedback.

**Figure 8.**
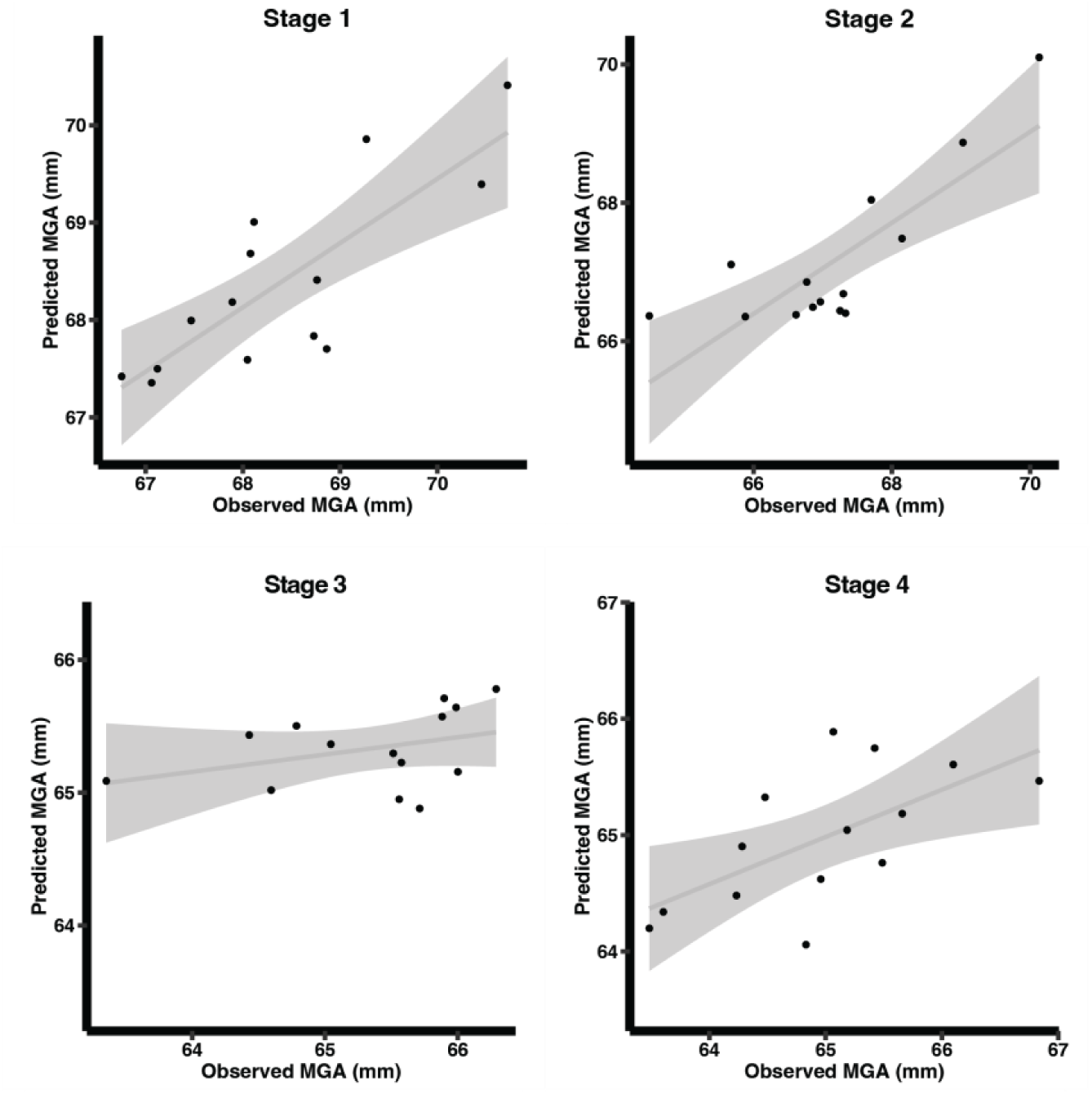
The correlation between the observed MGAs and the MGAs predicted by the models. The exponential models were used for Stages 1 and 2, while the linear models were employed for Stages 3 and 4. All stages, except for Stage 3, showed significant correlations. The gray area denotes 95% between-subject confidence interval.

Introducing additional depth cues to the stimulus in Stage 2 (multi-cue stimulus with same physical ground truth as the disparity only stimulus in Stage 1) led to an abrupt increase in MGA, followed by a trial-to-trial decrease in grip apertures based on the haptic feedback, consistent with a further overestimation bias. The exponential model (*AIC* = 40.9) explained this trend in MGAs better than the constant model (*AIC* = 51.93) and the correlation between the observed MGAs and the MGAs predicted by exponential model was again significant, *r*(12)= .81*, p < .*001 (Figure 8). These results indicate that the inclusion of more depth cues resulted in a greater overestimation of depth than the single-cue stimulus, causing greater planned grip apertures which rapidly adapted based on the physically correct haptic feedback. The observed patterns of MGAs in the first two stages were consistent with the prediction that there are perceptual and visuo-motor biases in the perception of depth (Figure 4).

In Stage 3, however, we did not observe any systematic changes in grip aperture trial-to-trial, with the constant model (*AIC* = 36.52) exhibiting comparable fits to the linear model (model assuming the presence of an aftereffect; *AIC* = 36.56). Moreover, the MGAs predicted by the linear (aftereffect) model were not significantly correlated with the observed MGAs, *r*(12)= .36*, p* = .205 (Figure 8). This suggests that there was no aftereffect as would be predicted by prior visuomotor adaptation observed in Stage 2.

Finally, in Stage 4, the MGAs tended to decrease across bins, as indicated by the better fit of the linear model (*AIC* = 35.2*,slope* =−.14) compared to the constant model (*AIC* = 40.49) to the data. The linear model’s predicted MGAs were also significantly correlated with the actual MGAs, *r*(12)= .64*, p* = .014 (Figure 8). This decreasing pattern is unexpected because the perceived depth of the multi-cue stimulus was matched to the haptically-specified depth, which should not trigger any adaptation. Even if the overestimation bias did not exist and the object appeared shallower as its rendered depth decreased, the planned grip should have shown a sudden decline and then increased back to the haptically-specified depth, which is the opposite of what we observed.

Overall, across the 4 stages, while the first two stages are consistent with an overestimation bias for both single and multi-cue stimuli, the data from Stage 3 were not consistent with an post-adaptation aftereffect, and the data from Stage 4 unexpectedly revealed a linear decreasing trend inconsistent with any plausible models of bias in perception and action (compare to Figures 4 and 5). However, inspecting the overall pattern of MGA revealed a general decreasing trend in MGAs across all four stages. This pattern is consistent with a practice effect on grasp safety margin that is often observed in experimental measurement of MGAs. The safety margin is a marginal space the motor system supplements to the object size or depth estimate to prevent collisions between fingertips and the object^57,58^. The safety margin tends to increase with higher uncertainty in action, such as reaching a distant object quickly or the hand being invisible, but then reduces with practice^17,59–62^. In our experiment, participants were required to perform rapid and natural grasps. Notably, to prevent the effect of visual feedback on grasp trajectories, neither the hand or index finger was visible during the grasp. This would cause a high level of uncertainty at the beginning of each stage which may increase the safety margin. This uncertainty reduces, along with the safety margin, with trial-to-trial practice, which could explain an overall trend of decreasing MGA across the entire experiment. If this was the case, this trend potentially exaggerated the adaptation effect in Stages 1 and 2, nullified the aftereffect in Stage 3, and caused an inexplicable decreasing pattern in the MGA in Stage 4. To address this practice effect, which might be obscuring the genuine adaptation and aftereffect effects across all stages, we detrended the data to remove this confounding practice-related trend in MGAs. Specifically, we discounted the linear trend by subtracting the least-squares fit linear function from each participant’s entire MGA data set across the 4 stages.

The detrended MGA revealed a clearer pattern of planned grip (Figure 9), consistent with the prediction of a sensory and visuomotor bias (Figure 4). In Stage 1, the single-cue stimulus was still overestimated as the visuomotor adaptation was evident following an exponential decay function (*AIC* = 35.7; the constant model did not show comparable fit: *AIC* = 40.46). The physically-matched multi-cue stimuli also yielded overestimation in planned grip (exponential model fit: *AIC* = 39.53) which was better than the constant model (*AIC* = 46.39). For both Stages 1 and 2, MGAs predicted by the exponential model again showed significant correlations with the actual MGAs, *r*(12)= .68*, p* = .007 and *r*(12)= .73*, p* = .003, respectively (Figure 10). It is noteworthy that the overestimation persisted in both Stages 1 and 2 even after removing the potential practice effect, since detrending would attenuate the exponential decay pattern observed in the original MGA dataset. This indicates that the sudden increase in MGA followed by adaptation in these stages is primarily attributable to an inherent overestimation bias in depth perception, rather than being influenced by other confounding biomechanical factors.

**Figure 9.**
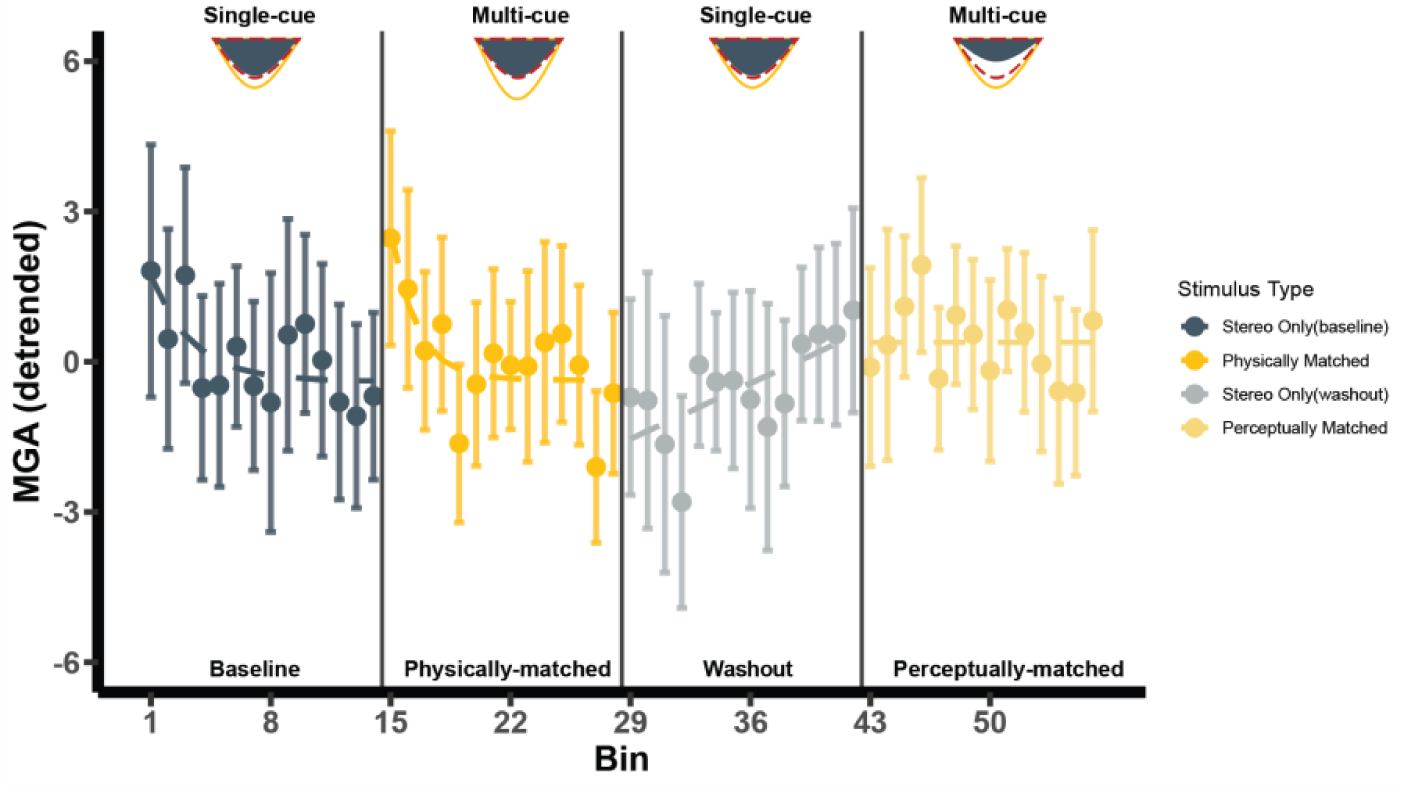
Grasping task results after discounting the linear trend attributed to practice effects. Each bin consists of 3 trials of varying depth magnitudes plotted on the x-axis, while the average detrended MGA of each bin is plotted on the y-axis. The dashed line is the best-fitting model: exponential models for Stages 1 and 2, the linear model for Stage 3, and the constant model for Stage 4. The error bar denotes 95% within-subject confidence intervals^54^.

**Figure 10.**
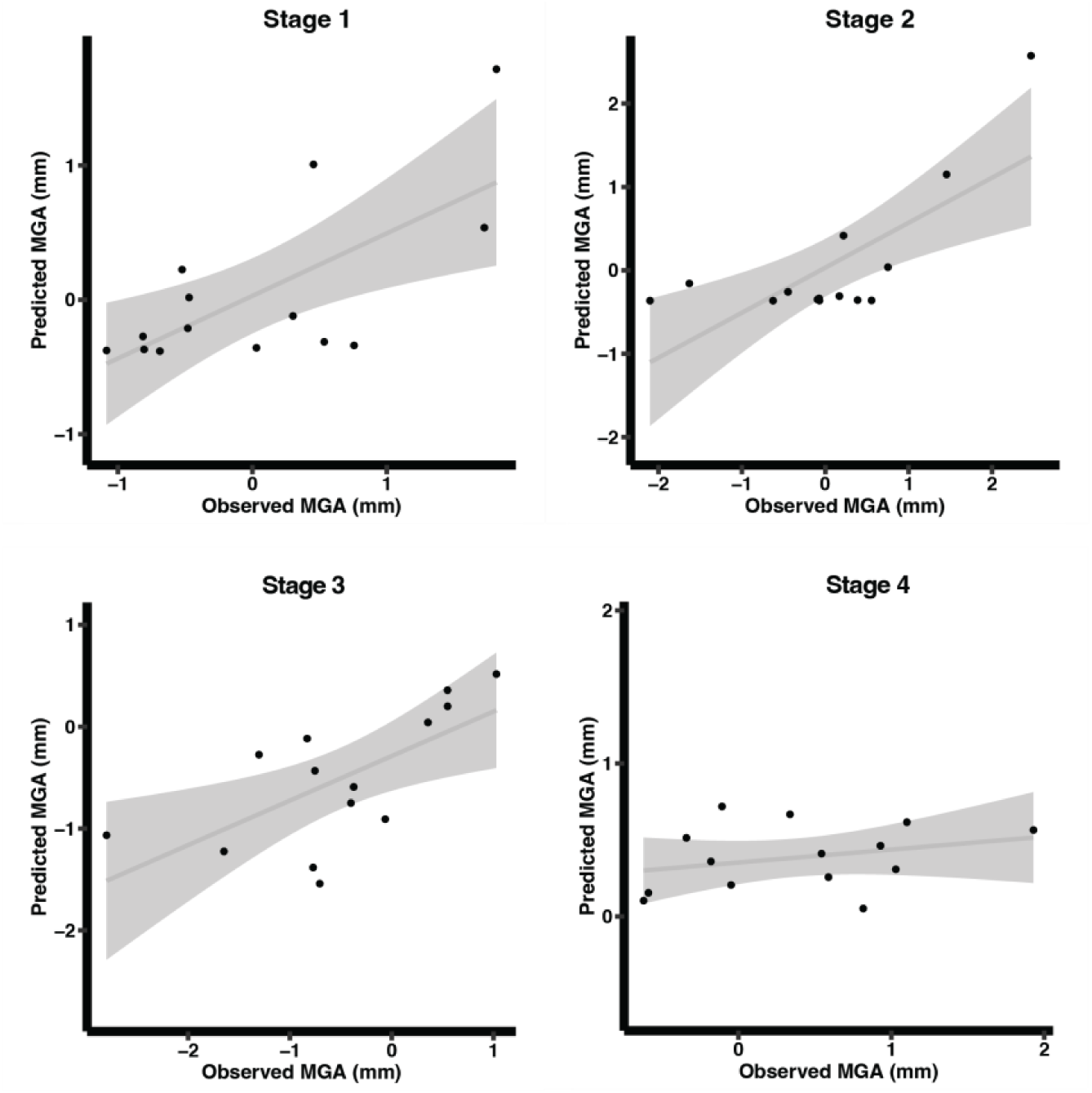
The correlation between the detrended MGAs and the MGAs predicted by the models. For Stages 1 and 2, the exponential models’ predicted MGAs were significantly correlated with the detrended MGAs. The predicted MGAs from the linear models were significantly correlated with the MGAs in Stage 3, but not in Stage 4. The gray area denotes 95% between-subject confidence interval.

When the stimulus switched back to the single-cue stimulus in Stage 3, the MGA was reduced in the beginning consistent with an aftereffect but converged back to the baseline after receiving haptic feedback, evidenced by a better fit of the linear model (*AIC* = 36.67) than the constant model (*AIC* = 42.72). Also, the observed MGAs were significantly correlated with the MGAs predicted by the linear model, *r*(12)= .66*, p* = .01 (Figure 10). In Stage 4, where the multi-cue stimulus was perceptually matched to the single cue stimulus of Stage 3 there was no change in planned grip (constant model: *AIC* = 34.21, exponential model *AIC* = 37.68). Although the linear model showed a comparable fit (*AIC* = 34.97) to the constant model, its slope was negative (−0.05) indicating that the MGA decreased across bins, which does not align with either prediction (Figures 4 and 5). Additionally, the MGAs predicted by the linear model were not correlated with the observed MGAs, *r*(12)= .29*, p* = .314. These results suggest that the visuomotor system did not detect any discrepancy between the perceived depths of single-cue stimulus and perceptually matched multi-cue stimulus, although their rendered depths were different. The overall pattern of results in the practice-detrended data is clearly consistent with the prediction of an overestimation bias in disparity-only stimuli and a further overestimation in multi-cue stimuli within reach space (Figure 4).

## Discussion

It has been widely accepted that error-free visually guided actions result from accurate visual estimation of spatial parameters for perception or action, despite abundant contradicting empirical evidence. To address this controversy, we capitalized on visuomotor adaptation as a hallmark indicator of bias in perceptual estimates. In a preliminary perceptual matching test, we replicated previous findings that objects appear deeper when specified by multiple cues than by a single cue. In the main grasping experiment, we found clear evidence of overestimation bias in both perceptual and visuomotor estimates within reach space. Specifically, there was overestimation in planned grip and adaptation to haptic feedback for the single-cue object, which aligns with previous studies showing overestimation from the disparity cue alone within reaching distance^10–12^. Crucially, further overestimation and adaptation effects were found for the multi-cue object having the same rendered depth as the single-cue object, indicating different perceived depths. However, no such effect was evident for a multi-cue stimulus which had a different rendered depth adjusted to match the perceived depth of the single-cue stimulus. These findings confirm the presence of systematic biases in visual estimates for both perception and action.

The present findings contradict a central claim of two mainstream views: veridical perception and veridical action. The veridical perception view proposes that the sensory system achieves veridical perception by optimally integrating information from various sensory cues that are themselves inherently unbiased^20–30^. Some proponents suggest that so-called flatness cues could explain estimation bias caused by increasing the number of depth cues^16,31^. In virtual viewing, depth cues like accommodation, the retinal blur gradient, and sometimes disparity, specify the zero depth of the flat screen, putatively weakening the effect of cues specifying non-zero depth. Flatness cues would be less influential as more non-zero depth cues are introduced, potentially explaining why multi-cue stimuli appear deeper than single-cue stimuli. However, recent studies showed that the presence of flatness cues does not contribute to the magnitude of perceived depth. In a wide variety of studies, perceived depth remained unaffected in the presence or absence of a variety of flatness cues including visible frame of the display, focus, screen pixelation, and disparity, specified a flat surface^18,19,63–66^. Moreover, in our findings (Figures 9 and 10), if flatness cues had an impact, single-cue stimuli should have appeared shallower than physically specified, resulting in an underestimation of planned grip, which is the opposite of what was observed in our experiment, Stage 1.

Another prominent view, the veridical action view posits that while sensory estimates for perception may be biased, those guiding actions nevertheless remain veridical because they are derived from a processing stream that is distinct from that underlying perception^33–35^. According to this view, the grip size should be accurately determined by the rendered physcial depth, regardless of any bias in perceived depth. Thus, in the current study, the planned grips in the first three stages, where the rendered depth was the same, should have remained constant, with a sudden decrease only in Stage 4 when the rendered depth was reduced (Figure 5). Contrary to this prediction, we observed that the planned grip was influenced by the perceived depth, which varied with the number of depth cues. The planned grip only stabilized around the haptically specified depth after the visuomotor adaptation corrected the error. This also suggests that previous studies may not have detected errors in action because the errors were masked by averaging MGAs across multiple trials. Since visuomotor adaptation rapidly reduces error based on the sensory feedback, even initially biased grasps would become accurate within a few repetitions, resulting in accurate planned grips in most trials.

There are, however, some aspects of the data visible on closer inspection that need to be addressed when making the above interpretations and conclusions. In Stage 3, we found a significant linear increase in MGA (after detrending) as expected by a re-adaptation effect following a negative aftereffect. However, one could question why the average MGAs across the first two bins of Stage 3 do not seem smaller than the last few bins of Stage 2. One explanation is that the MGA measure is affected by biomechanical noise which can cause random variations that make it difficult to directly compare one data point to another. A second possible reason is the tendency to maintain stable movement patterns based on prior experience^67–69^. Movement planning relies on both current sensory signals and expectations formed from prior experience. Repeated exposure to similar signals improves precision by reducing movement variability but can reduce accuracy when sudden changes occur. In late Stage 2, the visuomotor system may have developed an expectation of stable signals from the multi-cue stimulus. This expectation likely carried into early Stage 3, so reductions in visual depth estimates may not have immediately affected MGA. However, with repeated exposure to reduced depth, the visuomotor system may have updated its expectations and adjusted MGA based on the signals from the single-cue stimulus.

Another possible reason could be the use of multiple depth magnitudes. Visuomotor adaptation occurs by comparing the expected and actual outcomes of an action and compensating for the discrepancy between them. The aftereffect subsequent to visuomotor adaptation is more obvious and less noisy when that discrepancy has a constant value (e.g., grasping the same object repeatedly)^70^. However, in our experiment, we instead test three depth magnitudes to prevent participants from employing a strategy to repeat a memorized grip aperture without visually estimating the object depth. Consequently, the magnitude of the discrepancy between the depth estimate (both visual and visuomotor) and the depth specified by the haptic feedback varied depending on the object depth, which might have attenuated the measured aftereffect when comparing single data points.

It is important to note that the present study utilized a natural perceptual bias—the tendency to perceive greater depth with more depth cues—rather than artificially inducing conflicts between sensory signals. Previous studies typically induced visual illusions, conflicts between visual cues, or perturbations in sensory feedback^20,37,40,56,71^. These approaches allow independent manipulation of multiple sensory information, but such conflicts are rarely encountered in real life. In contrast, our grasping task featured consistent depth information from binocular disparity and pictorial cues and still revealed an overestimation bias, providing stronger evidence of the existence of inherent systematic biases in visual and visuomotor sensory estimates. Furthermore, our grasping task was designed to measure the planned grip automatically determined by the perceived depth. This implicit measure prevented conscious recognition or adjustment for biases, enabling the visuomotor system to detect their existence by itself. Our study highlights the potential use of visuomotor adaptation as a diagnostic tool for natural perceptual biases. While there is extensive literature on visuomotor adaptation, our study uniquely leveraged its role in detecting and compensating for errors stemming from intrinsic perceptual biases.

Our results pose an interesting question: If sensory biases like the ones confirmed here are widespread, how do we appear to successfully execute visually guided actions? Previous studies suggest that the visuomotor system compensates for errors through online control and adaptation. Constant online visual feedback, such as vision of the finger positions or the relative distance between fingers and objects, helps reduce possible errors and readjusts motor commands^39,41,57,72,73^. Moreover, sensory feedback from past experiences, like haptic feedback from touching objects, triggers visuomotor adaptation to improve subsequent actions, as discussed here and in related studies^56,70,74,75^. Consequently, these flexible adjustments based on sensory feedback, visual and visuomotor systems suggest that perceptual estimation of 3D properties are not always required to be veridical or accurate.

In conclusion, the results of this study challenge the prevailing assumptions about the veridicality of spatial perception and visuomotor performance. By leveraging visuomotor adaptation as a valuable tool to study bias in depth estimation, we demonstrate that perceptual biases do exist and significantly affect motor planning. Although this study used virtual stimuli to systematically manipulate the availability of different depth cues, future research should examine whether these findings generalize to more natural viewing conditions using physical objects, without being limited to virtual environments.

## Methods

### Participants

To estimate the minimum sample size, we conducted a power analysis by simulating data based on parameters from a prior study^76^ that examined visuomotor adaptation resulting from the overestimation of depth in multi-cue stimuli. The analysis indicated that a minimum of 11 participants would be required to detect the superior fit of the exponential model compared to a constant model with 80% statistical power. To account for potential variability due to task-related noise and biomechanical constraints, a larger sample was recruited. Twenty-seven participants from Brown University participated in the study. However, two participants were excluded because their finger positions were frequently unrecorded due to occlusions of finger-tracking sensors. Additionally, one participant whose planned grip (i.e., maximum grip aperture) deviated more than 2.5 standard deviations from the mean planned grip of all participants was identified as an outlier. Hence, twenty-four participants’ data were included in the further analyses. All participants were right-handed and had normal or corrected-to-normal vision. Participants gave informed consent before performing the task and received $12 per hour as compensation. The current study was approved by the Brown University Institutional Review Board (No. 0402991569) and conducted following the ethical standard set forth by the Declaration of Helsinki.

### Stimulus and apparatus

The experiment was conducted in a completely dark room with a custom-built virtual reality apparatus (Figure 3). Participants were instructed to place their chin on the chinrest and wear Neotek FE-1 active shutter glasses (NEOTEK, Pittsburgh, PA) synchronized with the refresh rate of the monitor screen (85 Hz) throughout the experiment. The stereoscopic images of the stimulus were presented with a 19-inch Sony GDM F520 CRT monitor (Sony, Tokyo, Japan) located to the left of the participants (Figure 3), using an NVIDIA RTX A5000 graphics card (NVIDIA Corporation, Santa Clara, CA). The monitor was physically positioned to account for the viewing distance (40cm), resulting in a negligible vergence-accommodation conflict. The rendered images were reflected by a half-silvered mirror slanted at 45 degrees to the frontoparallel plane, and thus they appeared to be a virtual 3D object floating in space. The back of the half-silvered mirror was covered by black cardboard, making the physical objects behind the mirror invisible to participants.

Each participant’s inter-ocular distance (IOD) was measured with a digital pupillometer (Reichert, Inc., Depew, NY) to personalize stereoscopic rendering by accurately positioning the cameras at each observer’s estimated nodal points. Shutter glasses then projected the left and right images to the respective eyes, creating a 3D experience tailored to each participant’s IOD. Since we are trying to identify perceptual biases in the context of a visuomotor task, it was important to validate the stereoscopic rendering regime, using individual IODs. To achieve this, after calibrating the alignment of the monitor and physical object, two expert observers with different IODs viewed the rendered stereoscopic images reflected in the half-silvered mirror while simultaneously observing the physical object to be grasped through the same mirror. Observers visually inspected and self-reported whether the virtual object precisely overlaid and coincided with the physical stimulus. Additionally, a virtual dot was rendered on the thumb marker indicating its position, allowing us to verify whether it aligned with the targeted location on the virtual object when the thumb contacted the corresponding point on the physical object. This procedure confirmed that the rendered stimulus’s location and depth accurately matched the physical object when viewed from the experimental position.

The stimulus had the shape of a Gaussian profile protruding toward the observer (Figure 1). The stimulus surface was colored red. The left and right sides of the stimulus were occluded by two black rectangular planes, so participants could not estimate the 3D shape from the contour of the stimulus. For the single-cue stimulus, binocular disparity was the only depth cue specifying depth. Even though the stimulus had a textured pattern on its surface, we disrupted the texture gradient by random rotations of the foreshortening axis of each texture element^77^. Thus, the texture pattern was not informative about depth. For the multi-cue stimulus, shading and texture gradient cues were added to the binocular disparity cue. Black polka dots were randomly placed on the surface, resulting in a texture gradient that was informative about depth. For both single-cue and multi-cue stimuli, the areas of polka dots covered approximately 85% of the object’s surface area. Previous studies have shown that binocular disparity plays a dominant role in perceived depth, particularly for stimuli within reaching distance^18,78^. This dominance may mask the potential effects of monocular cues in the multi-cue condition. To counter this effect, we blurred (blur kernel width = 8pxs, step size = 0.5pxs) the image presented to the non-dominant eye, reducing the precision of correspondence between images on the left and right retinae^38,79^. This adjustment weakens the disparity cue, making it more comparable to monocular cues and allowing us to observe their potential contribution in the multi-cue condition. Note that this manipulation should have no effect on the predictions of either the veridical perception or the veridical action view, as it simply reduces the reliability of the disparity signal, not its magnitude.

In the perceptual matching task, participants made responses by using a keyboard. In the grasping task, hand movement trajectory was recorded at a sampling rate of 85 Hz by Optotrak 3020 Certus motion capture system (NDI, Waterloo, Canada), and small sensors consisted of three infrared-emitting diodes that were attached to the thumb and index fingertips. Through calibration procedure, the 3D positions of fingertips were computed in real-time with sub-millimeter precision. We coupled a motion-capture system with a virtual rendering system, enabling us to display the real-time position of the thumb during each grasping action using a small stereoscopic sphere. To provide haptic feedback, we placed a physical object at an identical viewing distance from the observer. This haptic object was comprised of a set of 3D-printed boards attached to the front and back panels of a linear actuator (Phidgets Inc., Calgary, Canada). The outside face of the front board was aligned with the peak of the Gaussian-shaped virtual stimulus, while that of the back board was aligned with its base. In each trial, the linear actuator adjusted the gap between the two panels to match either the rendered or perceived depth of the virtual stimulus, depending on the stage. This adjustment was achieved by moving the back board along the depth axis while keeping the front board aligned with the peak of virtual stimulus. This setup allowed participants to receive haptic feedback while interacting with virtual stimuli. Customized C++ programs were used to control the stimulus and the linear actuator, and record responses from the keyboard or Optotrak. 3D virtual stimuli were rendered using OpenGL.

### Procedure

#### General Procedure

After providing informed consent and having their IOD measured, each participant performed a perceptual matching task and then a grasping task. The perceptual matching task involved generating a set of single-cue and multi-cue stimuli with the same perceived depths, personalized for each participant. The resulting perceptually-matched multi-cue stimulus set was used in Stage 4 of the grasping task. This perceptual matching task confirmed the previously observed bias that multi-cue stimuli appear deeper than single cue stimuli, and therefore that in order to perceptually match a disparity only stimulus, a multi-cue stimulus (disparity + texture + shading) need to have a smaller rendered depth. This was observed for all 24 participants. Mirror alignment, monitor position, calibration of the physical object providing haptic feedback, and finger-tracking markers were adjusted before each task.

#### Perceptual matching task

On each trial of the perceptual matching task, participants were presented with a pair of stimuli: one single-cue stimulus and one multi-cue stimulus. The task was to adjust the depth of the multi-cue stimulus until it appeared to have the same depth as the single-cue stimulus using a keyboard. Participants could view only one stimulus at a time but were allowed to switch between the two stimuli multiple times to make fine adjustments. The depth of the single-cue stimulus was held constant at either 27 mm, 30 mm, or 33 mm, and each of these three depth magnitudes was repeated ten times in a pseudo-randomized order, resulting in 30 trials in total. The perceptual matching task yielded three pairs of single-cue and multi-cue stimuli with matching perceived depths but differing rendered depths. These pairs would be used in the subsequent grasping task. Participants completed approximately 15 practice trials before the actual task.

#### Grasping task

During the grasping task, participants were instructed to reach and grasp a virtual object as naturally as possible. Each trial started with a virtual target stimulus presented after participants’ fingers remained at the designated home position. Haptic feedback was provided by placing the physical object at the location of the virtual stimulus. Importantly, precise alignment between the virtual and physical objects was ensured through a separate validation process. The grasping task comprised four stages, each associated with specific types of stimuli (Figure. 4). In Stages 1 and 3, a single-cue stimulus was presented with three depth magnitudes: small (27 mm), medium (30 mm), and large (33 mm). In Stage 2, a multi-cue stimulus with the same rendered depth as the single-cue stimuli appeared. In Stage 4, we presented a multi-cue stimulus that matched the perceived depth of the single-cue stimulus, which was determined based on the results of the perceptual matching task. In all stages, three depth magnitudes were shown in a pseudo-random order, ensuring that no depth magnitude repeated until the other two had been presented. The depth specified by the physical object always matched to the rendered depth of the single-cue stimulus throughout the grasping task. After 25 practice trials, participants completed a total of 168 trials, with 42 trials at each stage.

### Data Analyses

#### Perceptual matching task

In order to confirm that participants perceived multi-cue stimuli as deeper than single-cue stimuli, we tested whether the multi-cue stimulus depth adjusted by participants was significantly smaller than the corresponding single-cue stimulus depth. Paired t-tests were conducted on the conditional means, with each mean calculated as the average of each participant’s responses across the repeated trials. First, we averaged across three depth magnitudes for single-cue and multi-cue stimuli and compared their average depths. In addition, we tested whether the difference between single-cue and multi-cue stimuli is significant for each level of depth magnitudes by using paired t-tests with Bonferroni corrections.

#### Grasping task

The trajectories of thumb and index finger movement were sampled at 85 Hz and analyzed using custom R scripts. In the preprocessing of raw trajectory data, missing frames due to the occlusion of sensors were linearly interpolated. Then, the trajectory data was smoothed with a 20-Hz low-pass filter. The grip aperture was computed as a vector distance between the thumb and index fingertips. Trials were excluded from the analyses by the following criteria: 1) the proportion of missing frames exceeded 15% of the total number of frames, 2) the time elapsed between the movement onset and completion of action exceeded 1,500 ms, 3) the planned grip (i.e., maximum grip aperture) deviated more than 2.5 standard deviations from each participant’s conditional mean planned grip. Approximately 3.2% of the trials were excluded by these criteria.

We defined three trials of three different depth magnitudes as a bin, and averaged MGA for each bin. If estimation bias exists in both perception and action, we expect visuomotor adaptation in the first three stages but no adaptation in the last stage (Figure 4). As established by previous visuomotor adaptation literature, which conventionally demonstrated either exponential or linear decreases in error^55,56^, we applied either exponential^80^ or linear models to the bin-averaged MGAs for each stage, selecting the model that provided the better fit. Then, we compared its performance with constant models (i.e., a linear model with an intercept only) to assess the occurrence of visuomotor adaptation. The comparison of model performance was based on the Akaike Information Criterion (AIC), taking into account both the goodness of fit and the complexity of the model. In addition, we conducted correlational analyses to test whether the fitted model could predict the observed MGAs.

## Notes

### Competing Interest Statement

The authors have declared no competing interest.

